# Accuracy of the Lotka-Volterra Model fails in strongly coupled microbial consumer-resource systems

**DOI:** 10.1101/2025.02.14.638304

**Authors:** Michael Mustri, Quqiming Duan, Samraat Pawar

## Abstract

The generalized Lotka-Volterra (GLV) model is a cornerstone in theoretical ecology for modeling the dynamics emerging from pairwise species interactions within complex ecological communities. The GLV is also increasingly being used to infer species interactions and predict dynamics from empirical data on microbial communities in particular. However, despite its widespread use, the accuracy of the GLV’s pairwise interaction structure in capturing the unseen dynamics of microbial consumer-resource interactions—arising from resource competition and metabolite exchanges—remains unclear. Here, we rigorously quantify how well the GLV can represent the dynamics of a general mathematical model that encapsulates key consumer-resource processes in microbial communities. We find that the GLV significantly misrepresents the feasibility, stability, and reactivity of microbial communities above a threshold, biologically feasible level of consumer-resource coupling because it omits higher-order nonlinear interactions. We show that the probability of the GLV making inaccurate predictions can be quantified by a simple, empirically accessible measure of timescale separation between consumers and resources. These insights advance understanding of the temporal dynamics in resource-mediated microbial interactions and provide a method to gauge the GLV’s reliability under various empirical and theoretical scenarios.

## Introduction

Microbial communities play an integral role in ecosystems across scales, ranging from their influence on the metabolism of individual organisms (Huttenhower et al., 2012), to whole-ecosystem biogeochemical cycling (Falkowski et al., 2008). The functioning of any microbial community emerges from the interactions among its constituent species or “strains”. Developing methods to quantify and predict the underlying mechanisms and dynamics of these interactions is arguably the new holy grail in both theoretical and applied ecology (Dutton et al., 2021; Hough et al., 2020; Hu et al., 2022; Huttenhower et al., 2012; Marsland et al., 2020*a*; Shetty et al., 2022; Silverstein et al., 2023). The rapid development of molecular and ‘omics technologies means that relatively high-frequency and taxonomically well-resolved data on microbial communities are burgeoning (Abbasian et al., 2018; Saifuddin et al., 2019; Woodcroft et al., 2018). These data provide information on the metabolic capacities (“traits”) of individual microbial cells and the interactions between them, offering a means of linking changes in the composition of genetic and functional traits in the community to its emergent phenotype dynamics (Freilich et al., 2011; Reed et al., 2014; Sanchez et al., 2023).

Real-world microbial communities are typically composed of complex interaction networks between trillions of cells, thousands of strains, and hundreds of species. This complexity poses a significant challenge to developing a general theory for microbial communities. To tackle this challenge, both inferential and mechanistic mathematical models of microbial communities are needed (Dedrick et al., 2023; Momeni et al., 2017; van den Berg et al., 2022; Weiss et al., 2016). Inferential models are designed to deduce interactions and their effect on the dynamics of species abundances from empirical time-series data, remaining agnostic to the mechanisms that shape those interactions (Barbier et al., 2018). That is, these studies infer direct pairwise interactions between species from time series data obtained from various empirical approaches, ranging from co-culture experiments (Friedman et al., 2017) to co-occurrence network snapshots reconstructed from gene sequencing, including metagenome assemblies (Marino et al., 2014). This simplicity is advantageous in predicting the dynamical outcome of large microbial communities, where tracking every interaction and resource is impractical, and understanding the underlying mechanisms is not the primary focus.

In contrast to inferential models, mechanistic models aim to predict community dynamics from first principles by incorporating information on metabolic and physiological traits alongside cellular processes that drive the growth of populations through species-species and species-environment interactions (Kreft et al., 2017; Marsland et al., 2020*a*; van den Berg et al., 2022).

Inferential and mechanistic approaches complement each other and should ideally agree when compared at relevant spatial and temporal scales for any given microbial community. However, whether inferential pairwise models can accurately capture the inherent complexity of microbial interactions—particularly when these are indirectly generated (e.g., through cross-feeding and higher-order interactions)—and the resulting dynamics remains uncertain (Dedrick et al., 2023; Diamant et al., 2023; Gralka et al., 2020; Letten and Stouffer, 2019; Momeni et al., 2017; Muscarella and O’Dwyer, 2020; O’Dwyer, 2018).

Currently, the most widely used inferential model is the generalized Lotka-Volterra system of ODEs (henceforth, the “GLVM”) (García et al., 2023; Hu et al., 2022; van den Berg et al., 2022; Vega and Gore, 2018). In the GLVM, each population has an intrinsic growth rate, and any pair interacts through two coefficients that quantify their reciprocal effects on each other’s growth and abundance. Advances in high-throughput sequencing have enabled the collection of time-series data on microbial abundances. The GLV model can be fitted to these data to estimate interaction coefficients, providing insights into the structure and dynamics of the community.

Beyond empirical applications, the GLVM has also been used in a large body of theoretical research to derive dynamical properties of microbial communities, including feasibility (existence of a non-trivial equilibrium) and stability to perturbations as well as addition or removal of species (Butler and O’Dwyer, 2018; Clegg and Pawar, 2024; Cui et al., 2021; Dougoud et al., 2018). However, the validity of these theoretical results too depends on how well the pairwise model approximates the underlying, mechanistic, consumer-resource dynamics as an “effective” representation (Marsland et al., 2020*b*; O’Dwyer, 2018). In particular, it remains unclear whether the pivotal assumption of time-scale separation between consumer and resource populations holds under a sufficiently wide range of scenarios (Marsland et al., 2020*b*; O’Dwyer, 2018).

Here we compare the dynamic behavior of a fairly general and widely used mechanistic model of microbial consumer-resource dynamics (MiCRM) (Goldford et al., 2018; Marsland et al., 2020*a,b*) and its effective Generalized Lotka-Volterra approximation (GLVA, mathematically the same as the GLVM) and quantify the concordance between them across a range of biologically meaningful scenarios. Specifically, we focus on the GLVA’s error rate, that is, how often and by how much the population dynamics from the approximated model deviates from the “true” consumer-resource dynamics (including feasibility), as well as local stability and reactivity of the equilibria. We identify parameter spaces of the MiCRM that result in greater disagreement between the models. With the goal of providing both empiricists and theoreticians with a basis for identifying sources of errors in their inferences and generalizations, we also provide a method for quantifying the degree of time-scale separation, which is the root cause of the GLVA’s failure.

## Methods

Our overall approach is to first derive the GLVA from the MiCRM for each particular parameterization of the MiCRM, and then numerically integrate each MiCRM-GLVA ODE system paired to compare their dynamical behaviors.

### The Microbial Consumer Resource Model

The Microbial Consumer Resource Model (Goldford et al., 2018; Marsland et al., 2020*a*) is given by (Table 1):

**Table 1:**
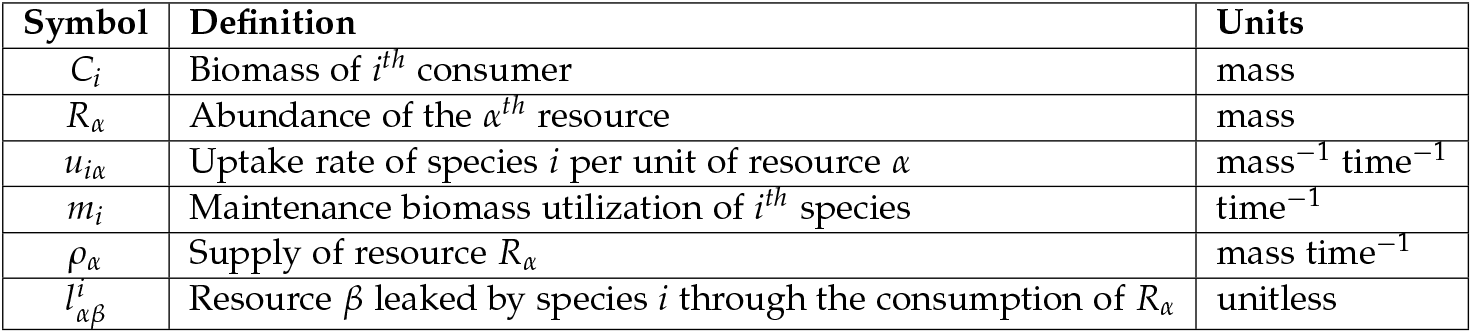
The MiCRM’s parameters.

| Symbol | Definition | Units |
| --- | --- | --- |
| $C_i$ | Biomass of $i^{th}$ consumer | mass |
| $R_\alpha$ | Abundance of the $\alpha^{th}$ resource | mass |
| $u_{i\alpha}$ | Uptake rate of species $i$ per unit of resource $\alpha$ | $\text{mass}^{-1} \text{time}^{-1}$ |
| $m_i$ | Maintenance biomass utilization of $i^{th}$ species | $\text{time}^{-1}$ |
| $\rho_\alpha$ | Supply of resource $R_\alpha$ | $\text{mass time}^{-1}$ |
| $l_{\alpha\beta}^i$ | Resource $\beta$ leaked by species $i$ through the consumption of $R_\alpha$ | unitless |

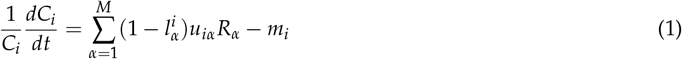

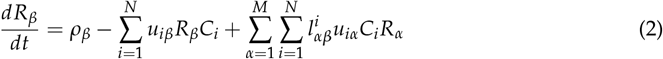

Here, Eqns 1 & 2 describe the time evolution of population (biomass) abundances of the *i*th consumer *C*_*i*_ and *α*th resource *R*_*α*_, respectively. For consumers, the growth rate is simply the sum of all resources consumed (*u*_*iα*_*R*_*β*_), offset by the proportion of resources leaked (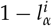, where 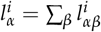 and a linear loss term representing respiration (*m*_*i*_). Resources are described by an arbitrary supply function (*ρ*_*β*_), the amount of resources taken up by all consumers (*u*_*iα*_*R*_*α*_*C*_*i*_), and the proportion of resources *R*_*α*_ leaked back into the system by consumers 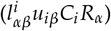.

Since the MiCRM is designed to depict two coupled interaction types: competition for shared resources and facilitation through metabolic cross-feeding, we expect the resulting system’s behavior to scale non-additively with respect to the number, magnitude, and type of interactions. This non-additivity stems from the dependence of interactions on resource abundances which, depending on community structure, may not be time-invariant. Therefore, given sufficient coupling (formally defined in Appendix C) for strength between the consumer and resource biomass pools, the dynamical properties of the community will be a nonlinear function of its size (number of consumers and resources) Pacheco et al. (2021).

### The Generalised Lotka-Volterra Approximation

The derivation of the GLVA is based on the approach originally used by MacArthur MacArthur (1970); Marsland et al. (2020*b*); first, assume resources reach equilibrium much faster than consumers, and then perform a first-order Taylor expansion around consumer equilibrium abundances Marsland et al. (2020*b*). This yields a resource-independent approximation to the MiCRM which can be rearranged to infer growth rates and pairwise interaction terms (derivation in Appendix A).

From the outset, the GLVA makes two strong assumptions: that interactions between populations are additive: the overall impact on a strain’s population in the presence of other strains is the sum of each of its pairwise interactions, and that the time scale of resource dynamics can be decoupled from those of consumers (Bosi et al., 2017; Momeni et al., 2017). As we show below, and as many others have noted beforehand (Butler and O’Dwyer, 2018; Momeni et al., 2017; O’Dwyer, 2018), neither can be guaranteed in complex microbial communities for a range of biologically realistic scenarios.

### Model parameterisation and simulations

To simplify the numerical simulations, maintenance biomass (*m*_*i*_) and resource supply (*ρ*_*α*_) were held equal and constant among all consumers and resources(*m*_*i*_ = *ρ*_*α*_ = 0.2). This choice of parameters assumes that consumers are energetically equivalent and comes with no loss of generality, since neither *m*_*i*_ nor *ρ*_*i*_ change the qualitative dynamics.

The consumer uptake matrix was drawn from a Dirichlet distribution defined by the consumer preference matrix *θ*_*iα*_ and a consumer specificity parameter Ω_*i*_. *θ*_*iα*_ represents the probability that consumer *i* will uptake resource *α* at a higher rate relative to other resources, while Ω_*i*_ defines how evenly the uptake is spread across the space of possible resources, with large values of Ω_*i*_ yielding generalist species and smaller ones yielding specialists.

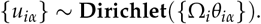

Using this particular method to generate the consumer uptake matrix allows us to fix its structure while including a degree of randomness. Furthermore, every uptake vector drawn from a Dirichlet distribution necessarily has a magnitude of 1, giving every consumer the same total uptake ability. If we wish to include scenarios where some consumers have a greater total uptake capacity than others, we can simply redefine the consumer uptake vectors as 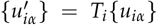, where *T*_*i*_ is the uptake capacity of consumer *i*. Here, however, we restrict the simulations to the case where all consumers have the same uptake capacity *T*_*i*_ = 1, the assumption being that added model complexity makes the MiCRM more difficult to approximate.

The consumer leakage tensor, which describes how consumers transform substrates into other resources, was generated analogously to the uptake matrix. Here, we must consider an additional parameter (*ϕ*_*iαβ*_) that encodes the probability of a substrate (*β*) being leaked by a given consumer (*i*) following the consumption of a particular resource (*α*). Hence, for every consumer-resource pair, we define a vector 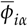 such that:

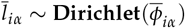

Further details about our sampling procedure are provided in Appendix D

### Quantifying Cross-Feeding and Niche Overlap

Consider a community with *N* consumers and *M* resources; the *N* × *M* matrix *u*_*iα*_ describes the space of preferences consumers have for resources. The vector 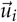 corresponds to the *i*^*th*^ row of *u*_*iα*_ and represents the distribution of resource preferences for the *i*^*th*^ consumer.

Recall our definition of Niche Overlap in the community as the average cosine similarity between each pair of consumer preference vectors 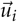:

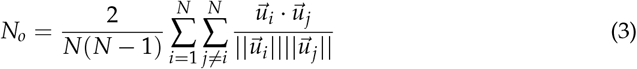

Indeed, this gives us a measure of how similar consumer preferences are on average, and since the consumer preference vectors are necessarily real-valued, it is a good scalar measure of preference overlap.

On the other hand, we may ask where consumer preferences concentrate (in resource space). Averaging the sum of the preference vectors easily accomplishes this.

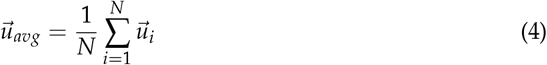

Notice that 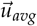 tells us very little about how preferences vary and, indeed, nothing of how they overlap. Coupled with *N*_*o*_, however, we can get a general notion of the shape defined by consumer preferences. While 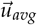 tells us the general direction of consumer preferences, *N*_*o*_ is a measure of the overall distance from that average. When *N*_*o*_ ≈ 1, we can surmise that consumer preferences are closely centered around 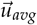, whereas if *N*_*o*_ ≈ 0, they are likely spread out.

To quantify how a given consumer is expected to leak resources in a complex community, we defined an effective leakage measure, 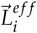, which is simply the overall sum of resource-specific leakage vectors, weighted by the consumer’s uptake capacity for the corresponding resources.

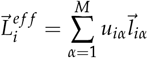

Where 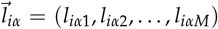 is the *i*^*th*^ consumer’s leakage vector for resource *R*_*α*_. This essentially measures the total distribution of metabolites leaked by a consumer in a scenario where every resource is available at equal concentrations.

Having defined effective leakage, we can measure the extent to which resources leaked by consumers will contribute to the community’s growth. To do this, we simply calculate the average pairwise cosine similarity between effective leakage and consumer uptake vectors.

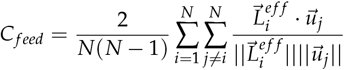

Therefore, *C*_*feed*_ is an overall measure of similarity between consumer uptake preferences and the distribution of resources that are likely to be leaked by consumers. Notice that we can reframe this cross-feeding measure in terms of average effective leakage, 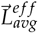.

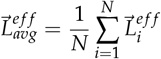

Hence we can consider *C*_*feed*_ to be somewhat dependent on the similarity between 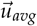 and 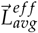.

More importantly, we can note that 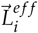 covaries with 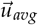 since they both depend on consumer uptake vectors 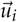.

### Stability and Reactivity

To evaluate the correspondence of stability and reactivity between the MiCRM and the GLVA, we calculated their respective Jacobian matrices at steady state, *A*_*Mi*_ and *A*_*LV*_, as well as the Hermitian parts of *A*_*Mi*_ and *A*_*LV*_ - *H*(*A*_*Mi*_) and *H*(*A*_*LV*_) Appendix B. Following this, stability characterised by the leading eigenvalues of the respective pairs of Jacobian matrices *λ*_*max*_(*A*_*Mi*_) and *λ*_*max*_(*A*_*LV*_), and reactivity the leading eigenvalues of the Hermitian parts *λ*_*max*_ [*H*(*A*_*Mi*_)] and *λ*_*max*_ [*H*(*A*_*LV*_)].

### Quantifying the GLVA’s Accuracy

To quantify the GLVA’s accuracy, we use the log-ratio of the GLVA- and MiCRM-predicted consumer abundances to obtain an instantaneous measure of error:

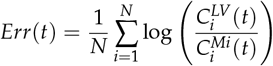

Using this quantity, we define both trajectory and equilibrium abundance errors. Trajectory error is measured by integrating *Err*(*t*) over the time it takes the system to equilibrate (*t*_*eq*_),

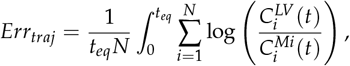

while equilibrium error is simply the instantaneous error rate at *t*_*eq*_:

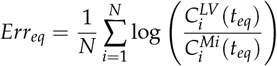

This method of calculating accuracy has the advantage of being insensitive to relatively small discrepancies and, at the same time, adequately capturing when there are more significant disagreements between models. This can be made obvious by noting that the error measure diverges when either model predicts an extinction that does not occur in the other.

We classified the GLVA’s accuracy by imposing a tolerance range of 20% above and below zero error at equilibrium; simulations within the range log(0.8) < *Err*_*eq*_ < log(1.2) were considered “accurate” whereas those exceeding the upper and lower limits were denoted “over-” and “undershot”, respectively.

## Results

### Cross-feeding Strength and Community Structure Influence GLVA Accuracy

To determine how the GLVA’s accuracy varies with metabolic leakage, we compared the proportion of GLVA simulations that remain within a 20% range below and above the objective MiCRM consumer abundances at equilibrium. Figure 2 illustrates how the GLVA goes from being accurate most of the time at low 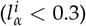, to failing roughly 50% of the time at intermediate 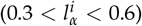, and failing consistently at high leakage 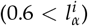. We note that most of the deviations in the predicted GLVA abundances stem from the fact that those particular communities converge on a subset of the consumer species richness relative to the corresponding “real” systems; that is, they do not reach the same composition (results not shown).

**Figure 1:**
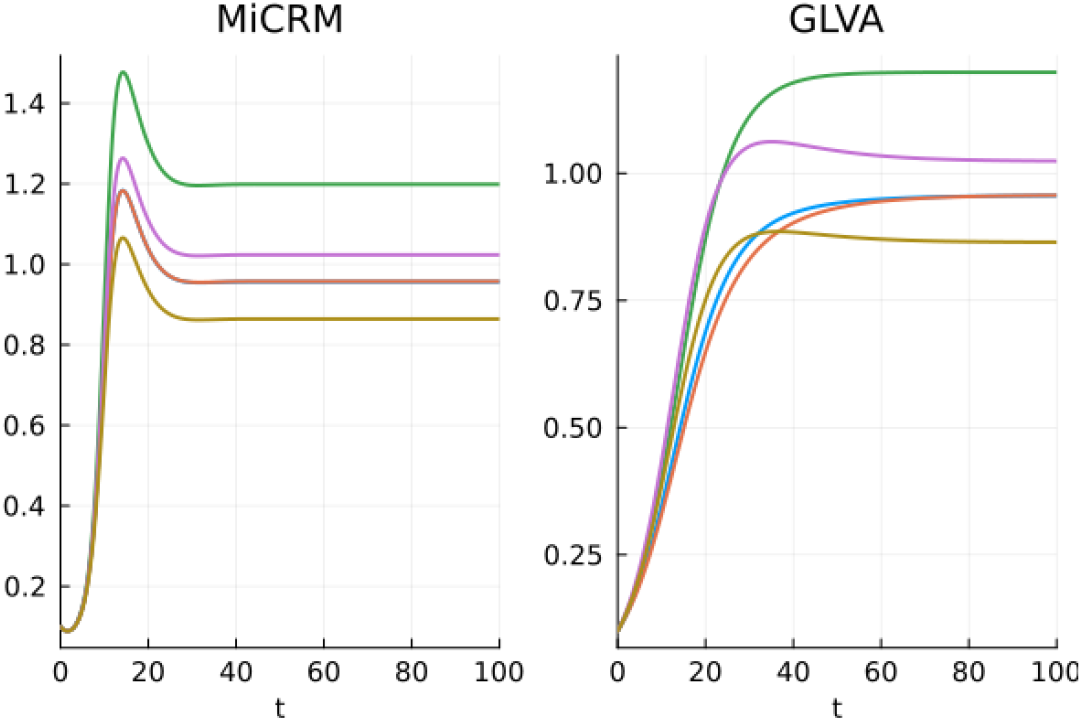
Example dynamics of the MiCRM and its corresponding GLVA. Plotted for a specific combination of parameters, demonstrating the differences in their transient dynamics.

**Figure 2:**
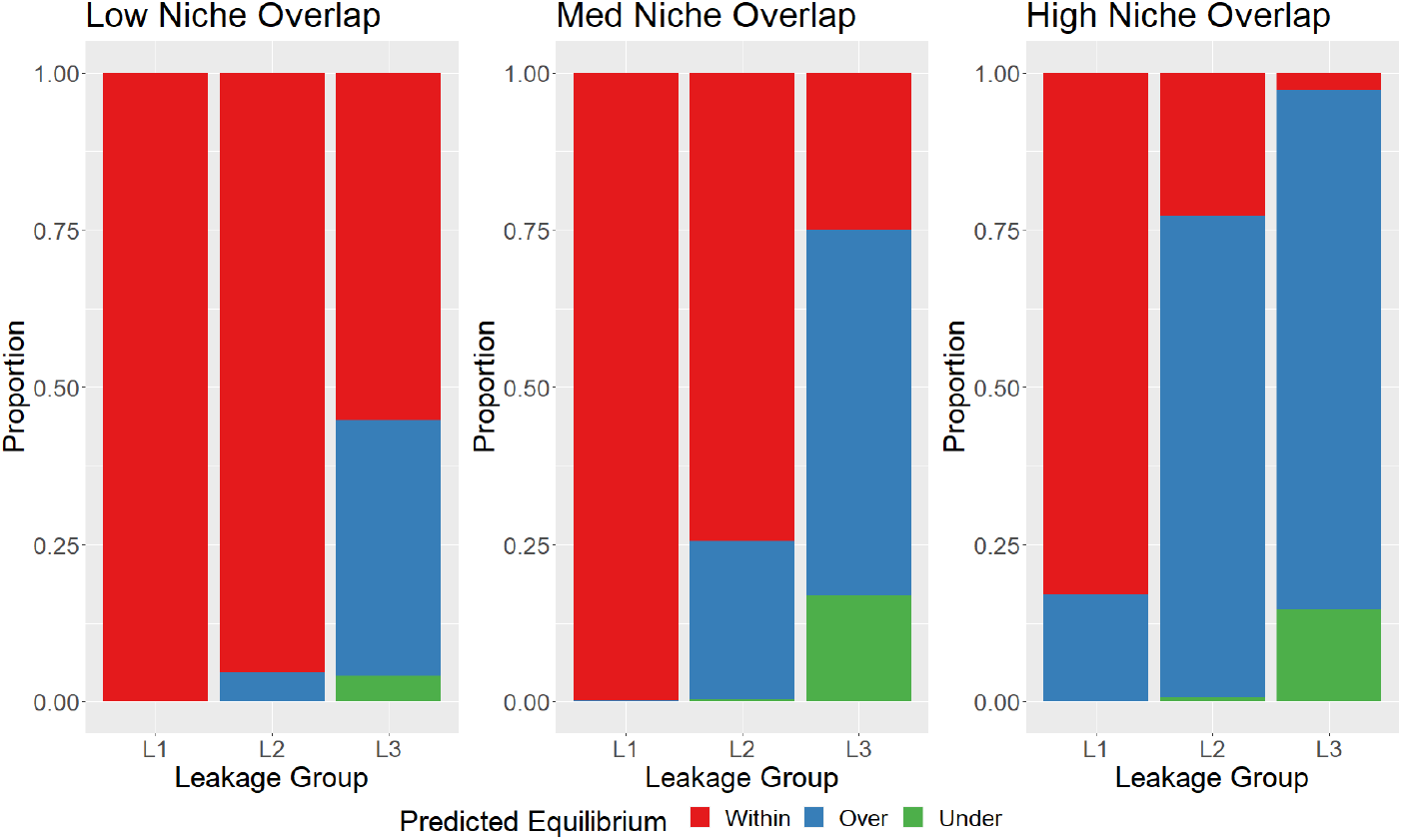
Effect of niche overlap and metabolite leakage on the GLVA’s accuracy. The proportion of simulations whose equilibrium abundances fell within (red), over (blue), or under (green) the 40% accuracy range for increasing levels of leakage given different degrees of niche overlap (0 ≤ *low* < 0.3, 0.3 ≤ *med* < 0.6, 0.6 ≤ *high*). Note that the effect of metabolite leakage, which governs the strength of coupling between consumers and resources, remains strong, largely irrespective of the niche overlap regime.

The biological relevance of such high leakage regimes in L3 is questionable, but it provides a useful limiting case. Additionally, we find that an interaction between leakage and niche overlap (NO) determines the GLVA’s accuracy (Fig.2), such that GLVA systems of highly competitive communities diverge more often at higher leakage values. On the other hand, we found no clear relationship between effective leakage stoichiometry (*C*_*feed*_) and GLVA accuracy.

### GLVA Underestimates Microbial Community Stability

While a simple comparison of trajectories and equilibria of the MiCRM and GLVA dynamics gives a picture of how well the GLVA can predict consumer abundances, it cannot predict their local stability and reactivity (Yang et al., 2023). That is, while both models may follow similar trajectories and settle on identical equilibria, this does not guarantee that they will respond to perturbations equivalently.

A direct comparison between the real parts of *λ*_*dom*_(*A*_*Mi*_) and *λ*_*dom*_(*A*_*LV*_), illustrated in Fig. 3 reveals some important discrepancies. The GLVA’s dominant eigenvalues typically have a less negative real part than the MiCRM (lower stability, i.e., longer return times after perturbations), with greater leakage pushing them closer to the imaginary axis, relative to the corresponding MiCRM dominant eigenvalues. That is, the GLVA, in general, predicts that communities are less stable than their MiCRM equivalents, with the discrepancy increasing with metabolite leakage.

**Figure 3:**
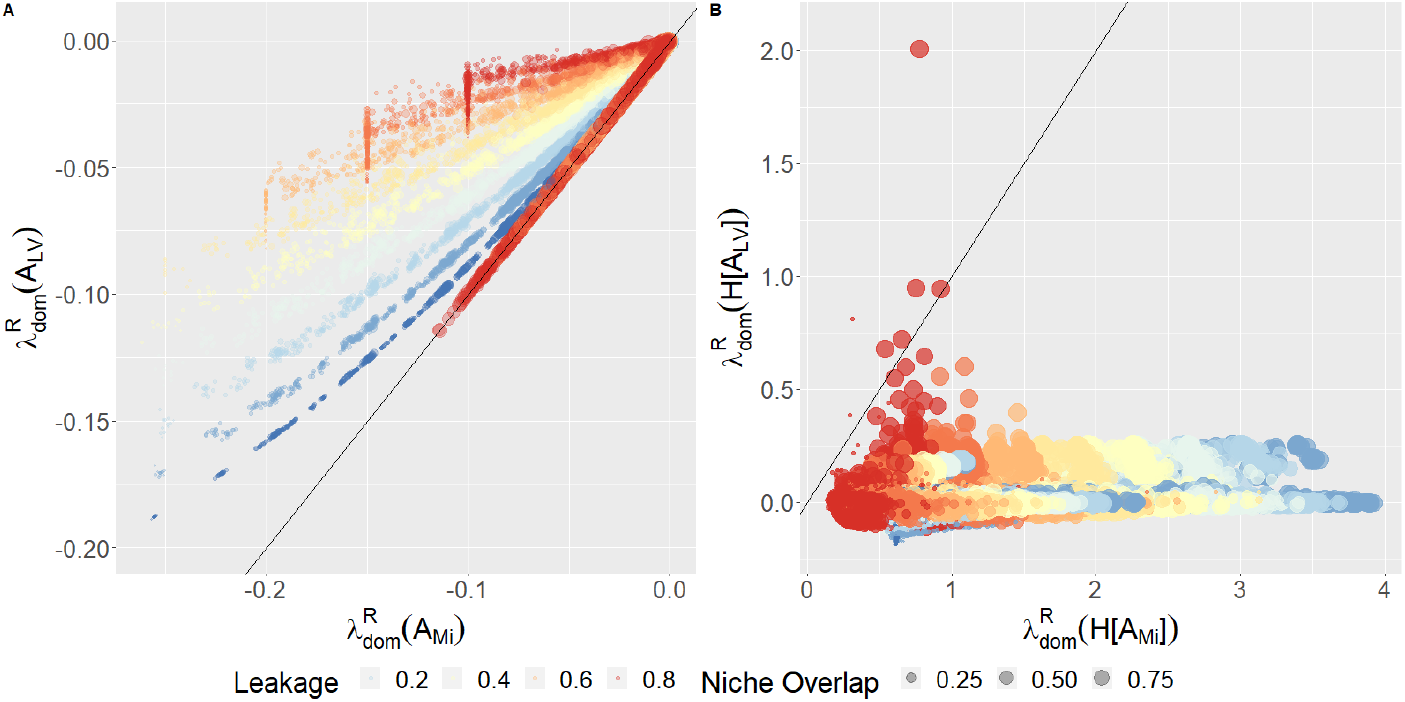
The GLVA underestimates both community stability and reactivity. The black lines represent 1:1 correspondence. All points lie on or above the black line in A, indicating lower-than-MiCRM stability, while all the points lying below the black line in B indicate higher-than-MiCRM reactivity.

Turning our attention to reactivity, we observe a starker disagreement between models. Whereas fixed points in the MiCRM are universally reactive, Re(*λ*_*dom*_ [*H*(*A*_*Mi*_)]) > 0, the same is not true for the GLVA. In fact, over half of the Hermitians evaluated in the GLVA had eigenvalues with negative real components. That is, these GLVA systems transiently dampened perturbations as opposed to the (ostensibly universal) amplification observed in the MiCRM.

### Consumer-Resource Coupling and the Causes of the GLVA’s Deviations

To better understand the GLVA’s inaccuracies, we need to consider two key issues: violation of fast resource dynamics assumption and non-additivity of interactions arising from significant higher-order terms in the consumer-resource dynamics. Evaluating additivity directly poses a particular challenge given the combinatorial nature of higher-order interactions. For example, a second-order analysis for a community of N consumers would require quantifying ∼*N*^3^ interaction coefficients between species triplets (see Guo and Boedicker (2016)). Even with simplifying symmetries and trivial pairs (i.e. *B*_*ijk*_ = *B*_*ikj*_ and *B*_*ijj*_ = *B*_*jji*_ = *B*_*jij*_ = 0), the number of coefficients would grow as *N*(*N* − 1)(*N* − 2). Fortunately, the time scales of resource dynamics, on the other hand, can be quantified rather straightforwardly as a method to predict the likelihood of the GLVA’s failure as we now show.

The validity of the assumption that resources reach equilibrium faster than consumers, which allows the time-scale separation for the approximation (Appendix A) depends upon the difference between “characteristic” (intrinsic) time scales. More precisely, the approximation will be within 𝒪(*ε*) of the real solution for times up to *t* ∼ 𝒪(*ε*^−1^), where *ε* represents the relationship between the slow (consumers) and fast (resources) time scales (Grimshaw, 2017; Strogatz, 2019), which can be quantified as:

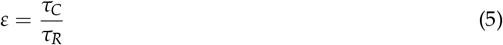

Here, *τ*_*C*_ and *τ*_*R*_ are the characteristic time scales of consumers and resources, respectively (further details in Appendix C).

To evaluate how the magnitude of *ε* affects the GLVA’s accuracy, we calculated the return times for consumers and resources directly from the MiCRM’s Jacobian matrix at steady state (Appendix C. We then compared the return time ratios between consumers and resources and chose the smallest one as a proxy for *ε*. Figure 4 shows that *ε* has a bimodal distribution, indicating an abrupt transition in relative time scales. In particular, simulations in which the GLVA had poor predictive power (fell outside of the equilibrium 80th percentiles) have significantly larger *ε*’s compared to cases which fell within, indicating failure of the fast resource dynamics assumption (timescale separation breaks down). However, looking at *ε* alone is not sufficient to confirm this. Instead, we must consider the separation of time scales relative to the evolution of the system as a whole. If the system reaches a stable equilibrium before exceeding the time frame that limits the approximation’s validity (*t* ∼ 𝒪 (*ε*^−1^)), we can expect the approximation to remain within a slim margin of error.

**Figure 4:**
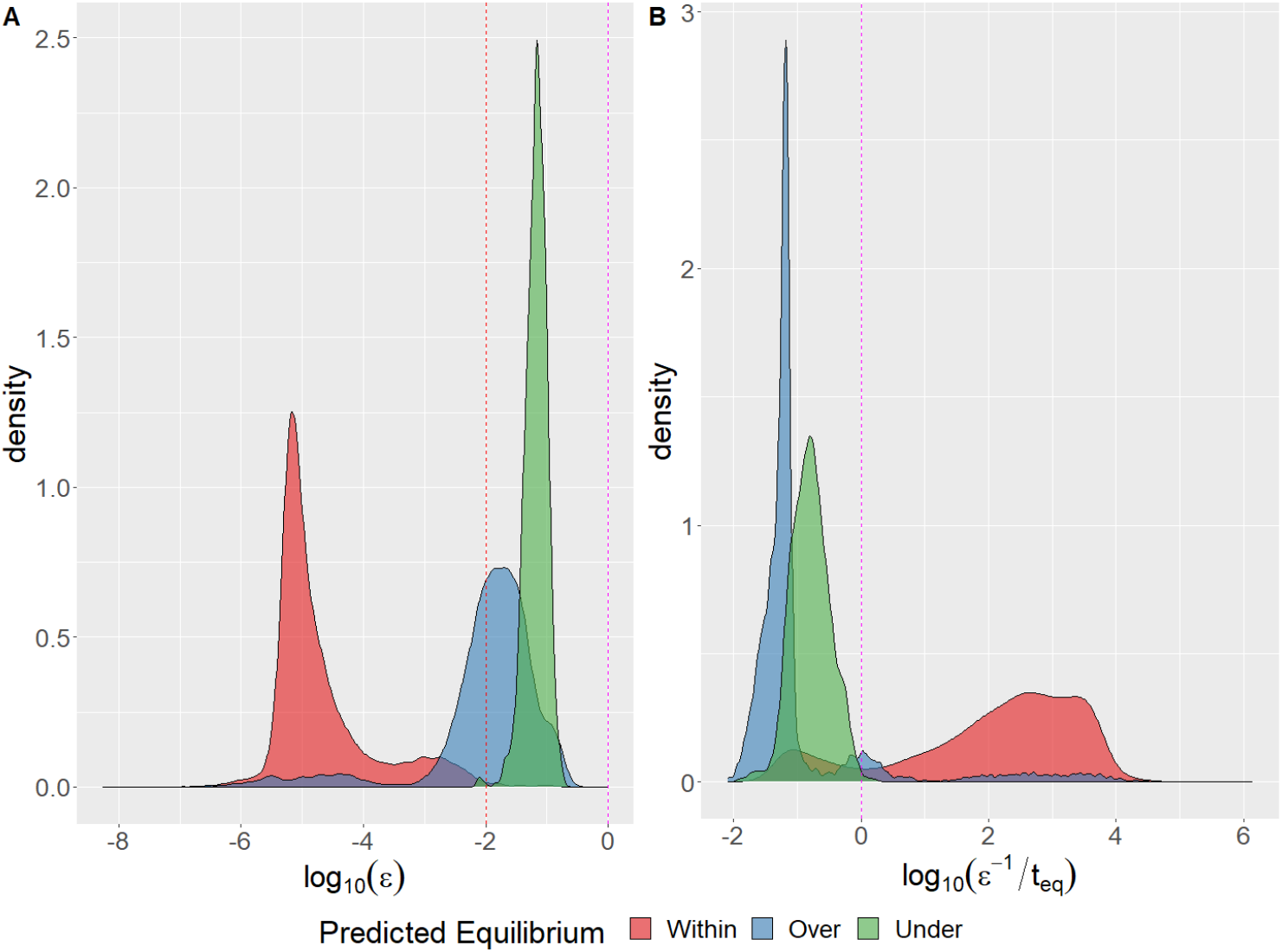
Effect of consumer-resource timescale separation on the GLVA’s accuracy. Distributions of return time ratios (*ε*) between consumers and resources (A) and *log*_10_(*ε*^−1^/*t*_*eq*_) (B) for simulations where the GLVA equilibrium abundances were within (red), over (blue), or under (green) the 20% tolerance range. Pink and red dotted lines correspond to *log*_10_(*x*) = 0 and *log*_10_(*x*) = −2, respectively.

We can confirm this last point by comparing *ε*^−1^ to the time to reach equilibrium in each simulation of the MiCRM. Figure 4B contrasts the distributions of log_10_(*ε*^−1^/*t*_*eq*_)—the magnitude of the timescale separation relative to the time to equilibrium—depending on where the GLVA’s equilibrium predictions fell relative to the accuracy threshold. The vast majority of simulations which fell outside this threshold have *ε*^−1^ << *t*_*eq*_, meaning that the span of time within which the approximation is valid is much smaller than the time to equilibrium. In other words, only if the time to equilibrium is shorter than *ε*^−1^ 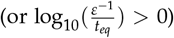, will the GLVA have a high likelihood of accurately predicting fixed points.

## Discussion

Although the generalized Lotka-Volterra model has been extensively studied theoretically and used empirically, its suitability for microbial communities in diverse settings remains unclear. By contrasting the dynamics of the MiCRM to its corresponding GLVA, we identified regions of parameter space where pairwise interaction terms did not accurately represent the underlying dynamics. In particular, we found that the accuracy of the GLVA (how well it predicts steady-state consumer abundances) declines rapidly with increasing metabolite leakage (and the resultant cross-feeding) and niche overlap. Furthermore, the GLVA’s dynamics qualitatively diverge from the underlying MiCRM: stability is particularly sensitive to leakage, while the deviation of reactivity from the true dynamics remains irrespective of the MiCRM’s parameterization. To the best of our knowledge, the accuracy of the GLVA’s dynamics in terms of reactivity has never been investigated. Our results reveal a potentially critical limitation of the GLVA because, as mentioned in Yang et al. (2023) and Hastings et al. (2018), intermittently perturbed reactive systems are more likely to settle into alternative stable states. Hence, if the approximation undermines the MiCRM’s response to perturbations, it no longer has any bearing on how the underlying system might shift under environmental or demographic stochasticity.

As mentioned in the Methods, our metric for GLVA’s accuracy focuses on discrepancies in relative abundances, meaning that it is highly sensitive to differences in the predicted survival of consumer populations at steady state. This implies that the increasing incidence of errors displayed in Figure 2 is likely due to mismatches in the equilibrium community composition (basically, the GLVA and its corresponding MiCRM systems settle on different stable states).

Interestingly, the GLVA asymmetrically errs on the side of coexistence, as seen from the predominance of simulations where the GLVA overshoots (Fig. 2). Only in high leakage scenarios (*L*3) is the converse more likely. Since pairwise interaction terms in the GLVA depend on resource concentrations at their steady state, this tendency to overshoot likely arises from the fact that the effective interactions differ both quantitatively (i.e., in terms of strength) and qualitatively (i.e., in terms of presence-absence or directionality) during the initial assembly stages, leading to the survival or extinction of strains before the community arrives at the interaction structure assumed by the GLVA.

The GLVA’s behavior at steady state differed from the MiCRM in two important respects: it systematically underestimates stability and fails to qualitatively capture the transient response to perturbations. The GLVA’s tendency towards less stable interaction structures (depicted in Fig. 3) reveals a key limitation of pairwise frameworks in that they disregard how resource concentrations constrain consumer populations. In effect, stability in the MiCRM directly incorporates the contributions of all resources, and while the GLVA is informed by resource concentrations at equilibrium, it cannot mimic the stabilizing effect of resource limitation. This likely explains why leakage, but not niche overlap, increases the stability gap between both models. Higher leakage leads to greater positive feedbacks between consumers, and in the absence of a compensatory response in resource concentrations, this has a net destabilizing effect in the GLVA.

While the MiCRM was reactive for every community simulated, the GLVA erroneously predicted the direction of the initial response to perturbations in over half of our data points. In the cases where the GLVA did display positive reactivity, it typically underestimated the strength of the effect by an order of magnitude. Though reactivity is not usually given as much attention as stability, it has been argued that it is at least as important for understanding the effects of chronically disturbed communities (Mari et al., 2017; Yang et al., 2023).

As shown in Fig. 4, the accuracy of the GLVA can be predicted from an approximate measure of the time-scale separation between consumers and resources. This apparent relationship between the temporal coupling of the consumer-resource system and the GLVA’s performance suggests that the degree of separation, as measured here, serves as a good indicator of when a particular consumer-resource system can be described with consumer abundances alone. More importantly, the degree of separation defines a neighborhood around equilibrium, within which the GLVA will remain below a bounded margin of error compared to the true dynamics. This neighborhood can be understood as the set of initial conditions (likewise, perturbations) for which *t*_*eq*_ ≤ *ε*^−1^.

It should be noted that this issue was originally raised, albeit indirectly, by MacArthur (1970) in terms of the symmetry of the matrix of interactions. If we take a Consumer-Resource model and assume that resources equilibrate quickly compared to the dynamics of the consumer, then the effective interaction matrix is necessarily symmetric. This result was later shown to be incorrect under a wider range of conditions, and as Marsland et al. (2020*b*) have shown, the requirement of symmetric interactions can be relaxed through certain rescaling procedures. More generally, it appears that some variants of the Consumer-Resource equations are inherently asymmetric and cannot be straightforwardly rescaled, making the symmetry of the interaction matrix an inadequate indicator of the GLV approximation’s validity.

While our analysis focuses on the GLVA’s ability to represent the dynamics emerging from metabolite competition and cross-feeding dynamics in the MiCRM, the results can be straightforwardly generalized to any interaction mechanism that is indirectly mediated, such as through chemical stressors (Smith et al., 2024). Indeed, our method of estimating *ε* (C) is directly applicable to any system, provided the ODEs can be defined, and the Jacobian is diagonally dominant.

As such, although the MiCRM’s structure is fairly general in its ability to capture the key elements of microbial consumer-resource dynamics, the question arises whether our results are applicable to variants of the MiCRM (e.g., different functional responses or the inclusion of essential nutrients). In particular, the issue of timescale separation and strength of consumer-resource coupling is a universal feature of coupled dynamical systems; hence, we do not expect any meaningful differences for a reasonably modified set of ODE’s ((O’Dwyer, 2018) addresses this point more directly). Whether these insights can be extended to accommodate more complex interaction mechanisms, such as allelopathy or chemically mediated signalling, remains to be seen.

Empirically determining *ε* in microbial communities requires precise monitoring of cell densities as well as substrate and metabolite concentrations. Given well-resolved data, *ε* can be estimated by perturbing the community and monitoring how long it takes the microbial consumer populations and substrate concentrations to return to a steady state. Another method would be to carry out single-strain culture experiments to determine each of the parameters (in this case: uptake, yield, and metabolite leakage), and use those data to calculate, using resource-explicit models like the MiCRM, the approximate timescale separation *a priori*.

Finally, we note that certain systems have intrinsic properties that allow a more robust timescale separation between the consumer and resource pools. Take the case of phytoplankton versus bacterial communities, for example. Phytoplankton cells, in general, are larger than bacteria and, therefore, operate at slower timescales relative to the metabolites they ingest and produce. Therefore, it is reasonable to assume that phytoplankton communities are only marginally affected by transient resource fluctuations, and the GLVA may, in general, be a more accurate representation of their dynamics.

Overall, these results provide guidelines for the appropriate use of pairwise models in the study of microbial communities. Firstly, theoreticians looking to use the generalized Lotka-Volterra model to relate community structure to the system’s dynamic behaviors should confirm that the system in question does not harbor environmental (resource) feedbacks strong enough to violate the model’s assumptions. This point becomes especially important when considering the impacts of environmental perturbations (e.g. temperature fluctuations (Bolnick et al., 2011; Clegg and Pawar, 2024; García et al., 2023; Smith et al., 2024; Vasseur et al., 2014)), a topic of increasing importance as we seek to quantify the effects of anthropogenic change.

A second point is that if we wish to understand how microbial interactions (both between species and with the environment) shape the assembly and function of the resulting communities, it is essential that empiricists and theoreticians focus on illuminating the mechanisms that drive those interactions. This includes developing detailed models that scale cellular and molecular processes (such as the effects of extracellular enzyme production, etc.) to population-level characteristics, as well as carrying out the necessary experimental studies with which to parameterize and validate such models. Emphasis should be placed on this latter point, given the numerous opportunities offered by the current state of “omic” technologies, live-cell imaging methods, and genome-scale metabolic modeling; it is now feasible to simulate microbial communities from first principles, as opposed to relying on *post hoc* reconstructions through statistical inference.

In conclusion, the magnitude of metabolite leakage and the resultant cross-feeding, mediated by niche overlap structure, has a strong bearing on the extent to which the GLVA can accurately reproduce the underlying MiCRM dynamics. In particular, we found near-zero correspondence between models with regard to reactivity, highlighting the importance of higher-order interactions and the GLVA’s inability to account for them in predicting perturbative behavior. We show that these inaccuracies stem primarily from (the lack of) timescale separation between consumer and resource dynamics, and provide a measure to quantify the strength of this coupling.

## A The Generalised Lotka-Volterra Approximation

Assuming a regime where resources achieve equilibrium much faster than consumers allows us to express the MiCRM dynamical equations solely in terms of the consumers and their interactions. Furthermore, we can reconfigure the terms in a way that gives us an equivalent, “effective” generalized Lotka-Volterra model.

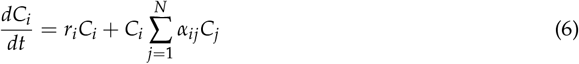

Fast resource equilibration implies that 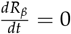, giving us the following relationship:

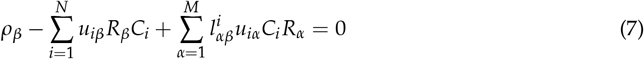

Solving for 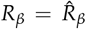, we can plug it back into the equation for consumers and perform a Taylor expansion of 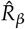 around the set of equilibrium populations *C*_*j*_ = *Ĉ*_*j*_.

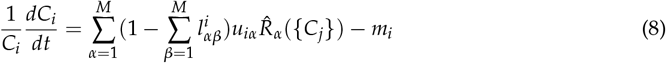

Expanding the Taylor series of 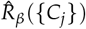 around equilibrium we have:

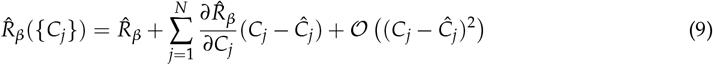

Our final expression for consumer dynamics is then

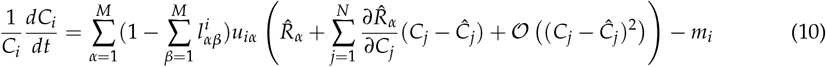

Here 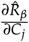 can be obtained through direct differentiation of 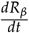, yielding:

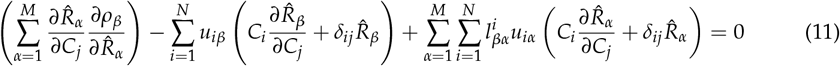

Solving the equation for the 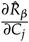 we obtain an expression in the form:

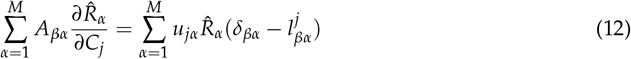

where

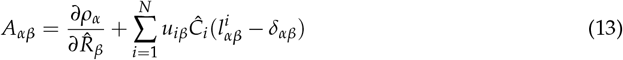

Inverting *A*_*αβ*_ (assuming it is an invertible matrix), we are able to obtain the partial derivatives explicitly.

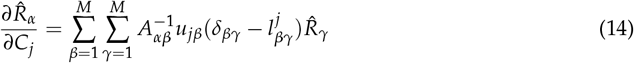

Finally, to obtain the effective Lotka-Volterra system, we must group terms and find the equivalent characteristic growth rates *r*_*i*_, and interaction coefficients *α*_*ij*_. After some algebra, we have the interaction matrix coefficients

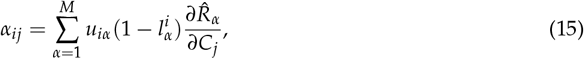

and their intrinsic growth rates,

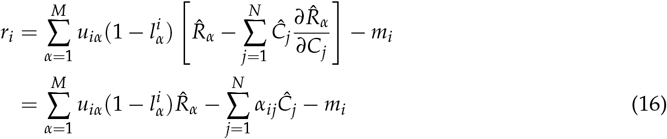

Having defined these coefficients, we can construct an equivalent effective GLV model for any given system of consumers and resources. The prerequisite is that our initial MiCRM system of equations must reach equilibrium asymptotically and that these equilibrium values (*Ĉ*_*i*_ and 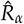) be known.

## B Stability Analysis

To quantify the characteristic timescales of both consumers and resources, as well as the stability of the feasible (equilibrium) microbial communities, we analyze the behavior of the linearized system about equilibrium.

We begin with trivial equilibrium solutions, 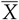 and 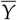:

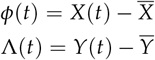

Next, we reevaluate them at a small perturbation away from the fixed points. For this, we differentiate them, which gives

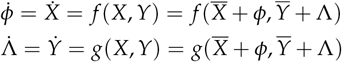

To simplify the analysis, we combine our dependent variables *X* and *Y* into a vector *Z*, and the response functions *f* (*X, Y*) and *g*(*X, Y*) into a single response function 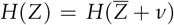, with *ν* our combined perturbation vector. We then perform a Taylor expansion around 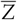

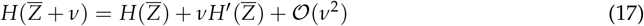

By definition 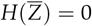, and since our perturbation is small (*ν* << 1) we neglect higher-order terms, thus

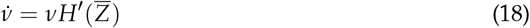

Where 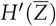 is the Jacobian of the response function evaluated at the fixed points 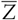. With this linearization, we can evaluate the behavior around equilibrium for any small perturbation *ν* given that 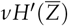 does not contain any non-trivial zeroes. Additionally, by performing an eigensolution decomposition of our linearized system, we can gain some insight into the stability of any non-hyperbolic fixed point (that is, fixed points for which the real parts of the eigenvalues of the Jacobian are non-zero).

To proceed, we must first explicitly derive the Jacobian of our system. We first note that the set of state variables {*Z*_*i*_} for *i* ∈ {1, …, *N*} is equivalent to {*C*_*i*_} and {*Z*_*i*_} = {*R*_*α*_} for *i* ∈ {*N* + 1, …, *N* + *M*}. Now, we can derive the diagonal terms for the consumer portion of the Jacobian (*i* = {1, …, *N*}).

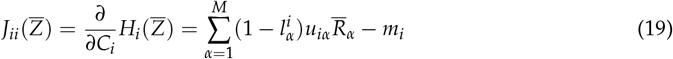

And the off-diagonal terms for the consumer portion are simply zero, *i, j* ∈ {1, …, *N*}:

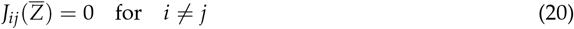

The off-diagonal terms for consumers with respect to resources are *α* ∈ {*N* + 1, …, *N* + *M*} *i* ∈ {1, …, *N*}:

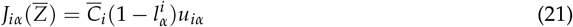

For the resource states, we can see that the diagonal terms will be *α* ∈ {*N* + 1, …, *N* + *M*}:

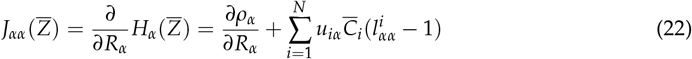

The off-diagonal terms with respect to consumers, *α* ∈ {*N* + 1, …, *N* + *M*} *i* ∈ {1, …, *N*}:

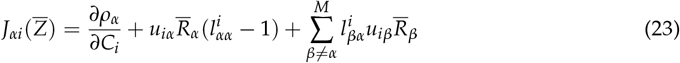

And finally, the off-diagonal terms with respect to resources, *α* ≠ *β α, β* ∈ {*N* + 1, …, *N* + *M*}:

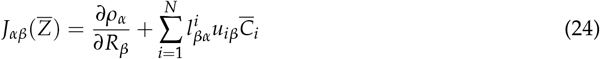

## C Quantifying separation of timescales (consumer-resource coupling)

To give a better intuition for the relevance of time-scale separation, we illustrate a simple example involving a driven harmonic oscillator with friction.

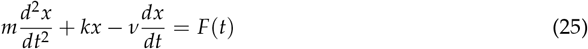

In the absence of forcing – *F*(*t*) = 0 – the solution is simply:

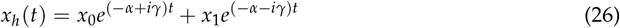

With 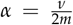 and 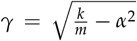. For an arbitrary *F*(*t*), the general solution can be obtained by applying a Laplace transform to both sides of equation 26.

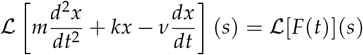

Employing the properties of the Laplace transform and re-arranging terms, we obtain the following relationship, where 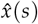 and 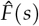 denote the Laplace transforms of *x*(*t*) and *F*(*t*), respectively:

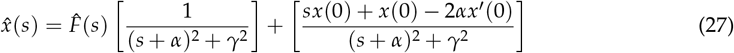

Taking the inverse transform of 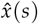, we note that *x*(*t*) is simply the solution to the damped harmonic oscillator (eq. 26) in addition to the convolution of *F*(*t*) with an exponentially decaying sine wave.

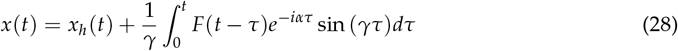

If we take *F*(*t*) to represent a coupling force between the oscillator and some external system (e.g. external resource supply), we can ask when *x*(*t*) can be treated independently of *F*(*t*)’s evolution. As is evident from equation 28, this is only possible when the convolution term (which we denote *F*_*_(*t*)) is vanishingly small. If we consider *F*(*t*) to be a periodic function or a decaying perturbation, then the question of whether *F*_*_(*t*) << *x*_*h*_(*t*) becomes a matter of relative timescales.

For example, if *F*(*t*) = cos(*ωt*), then the convolution becomes

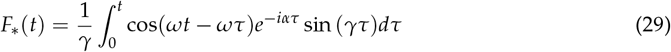

In general, *F*_*_(*t*) will decay to zero asymptotically and *x*(*t*) will converge to *x*_*h*_(*t*). However, unless *ω* >> *γ*, this decay will occur slowly, dominated by *α*. In other words, the deviation from *x*_*h*_(*t*) will persist until equilibrium unless *F*(*t*) evolves on a faster timescale. For a decaying perturbation such as *F*(*t*) = *a*_0_*e*^−*at*^, the argument is much the same. If *a* >> *α*, then *F*_*_(*t*) will quickly approach zero. Otherwise, *F*_*_(*t*)’s convergence will be dominated by *α*.

Before proceeding, it is important to note that, unlike the unilateral coupling function presented in this short example, consumer-resource coupling is bidirectional. That is, *F* is a function of *x*(*t*) and its derivatives, *F* = *F*(*t, x*^(*n*)^). This complicates matters significantly as deviations will no longer be bounded and it becomes quite possible for *x*(*t*) to assume a different trajectory.

Having defined the MiCRM’s Jacobian, it is possible to estimate the characteristic timescales of consumers and resources. Though this can be done rigorously by identifying the slow and fast manifolds, we present a simplified method that relies on the diagonal elements of the Jacobian. We first note that the decay rate following a small perturbation to either a single consumer or resource is given by:

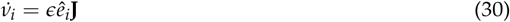

Here *ϵ* << 1 and *ê*_*i*_ is the basis vector for the *i*^th^ consumer or resource. It can be shown that return times following such a perturbation are inversely proportional to the corresponding diagonal element of the Jacobian.

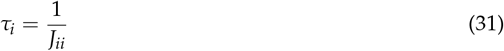

Hence, for consumers, we have

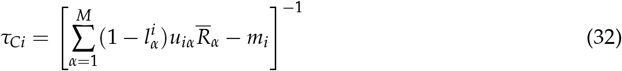

And for resources

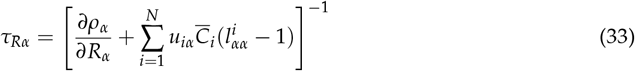

This is guaranteed so long as the Jacobian is diagonally dominant, which is generally true in the MiCRM, particularly for large systems. Next, we assume that consumer-resource coupling across the community is dominated by the fastest consumer and the slowest resource. We justify this assumption by noting that the presence of a single consumer-resource pair progressing at similar temporal scales can have cascading effects on the rest of the community; making it a sufficient condition for the violation of the fast resource dynamics assumption.

As such, finding the timescale ratio between consumers and resources is equivalent to:

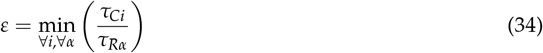

This is analogous to finding

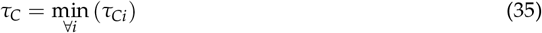

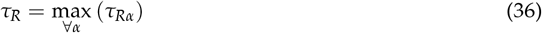

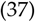

Hence:

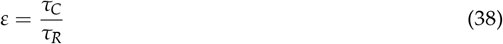

## D Balanced sampling of competitive and cooperative communities

To ensure that the community sampling procedure provides a well-represented range of competitive and cooperative structures, we designed the following method to vary parameters such that a desired average niche and effective leakage overlap is attained. We begin by constructing the consumer preference matrix. To this end, we define two extremes of the sampling procedure exemplified by a zero overlap matrix *θ*_*id*_, which is simply an identity matrix, and the maximum overlap matrix *θ*_*hom*_, which is a matrix populated by ones. The idea here is that we can define our overall consumer preference matrix as the weighted sum of the zero- and maximum-overlapping matrices; varying the weights in each extreme allows us to continuously move from zero to maximum overlap.

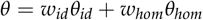

Having defined a method to continuously move between both edges of the overlap of consumer preferences, we now do the same with our specificity parameter Ω. The motivation behind this is that the specificity must be high if we want to reliably sample communities with negligible overlap. Conversely, as consumer preferences become more similar, we want specificity to be lower, allowing for greater variability and avoiding drawing the same highly homogeneous consumer uptake matrix multiple times.

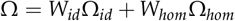

With this framework, we can define a niche similarity parameter *s*_*n*_ such that *w*_*id*_(*s*_*n*_) and *w*_*hom*_(*s*_*n*_) are decreasing and increasing functions of *s*_*n*_ respectively. Applying a similar argument to specificity, we need *W*_*id*_(*s*_*n*_) to be an increasing function and *W*_*hom*_(*s*_*n*_) to be decreasing. In short, we need the weights to be inversely proportional with respect to niche similarity, in this manner varying *s*_*n*_ between some arbitrary interval (say [0-1]), produces communities that go from no niche overlap to maximum niche overlap.

The stoichiometric matrix is sampled similarly to consumer uptake. However, the goal in this context is to create communities that exhibit a diverse array of cross-feeding behaviors. This is further complicated by the fact that cross-feeding is largely dependent on consumer preferences. To circumvent this problem, we sampled consumer leakage based on the maximum and minimum amounts of cross-feeding achievable for a given set of consumer preferences.

## Acknowledgments

We thank Alberto Pascual-García for insightful comments on an earlier version of this manuscript.

## Notes

### Competing Interest Statement

The authors have declared no competing interest.

